# scDblFinder in Python with GPU support

**DOI:** 10.64898/2026.08.12.744148

**Authors:** Andreas Hiropedi, Pierre-Luc Germain

## Abstract

High-throughput single-cell sequencing provides a scalable solution for characterizing cells and profiling gene expression for hundreds to millions of cells. However, this process gives rise to doublets, which can lead to inaccurate conclusions drawn from the data. A number of packages have therefore been developed to help accurately detect them, and in particular *scDblFinder* has been shown to outperform alternatives in the detection of doublets in single-cell (RNA) sequencing data. Being implemented in R, however, its adoption has been more limited in the Python community. Here, we present *scDblFinderPy*, a Python-based implementation of the *scDblFinder* R method, and show that it obtains similar performances. Furthermore, we include in it optional GPU support, thus further speeding up the process.

## Introduction

In single-cell RNA sequencing (scRNA-seq), doublets or multiplets are two or more cells captured within the same reaction volume (droplet or well) or (for combinatorial indexing) barcode combination, and sequenced together under a single cellular barcode. Doublets formed from different cell types (heterotypic doublets) can easily be misinterpreted as a distinct cell type and skew downstream analysis. Therefore, computational doublet-detection is a critical quality control step in single-cell workflows.

The *scDblFinder* R method (Germain et al., 2022) has been repeatedly and independently shown to outperform alternatives in the detection of doublets in single-cell (RNA) sequencing data (Germain et al., 2022; Heumos et al., 2023; Schriever and Kostka, 2023; Xi and Li, 2021a,b). Being implemented in R, however, has limited its adoption to the Python community, which often still operates with sub-standard solutions. In addition, while *scDblFinder* compared favourably with alternatives in terms of running time, the continuous increase in dataset sizes – especially within a single capture – has motivated the push for computational efficiency. To address these needs, we re-implemented the main *scDblFinder* functionalities in Python, also enabling optional GPU usage via the *rapids-singlecell* package (Dicks et al., 2026).

We benchmarked our Python implementation against the original R method, as well as against other prominent doublet-detection methods, using the benchmark datasets from Xi and Li, 2021b. We found that our Python implementation achieves comparable accuracy to the R package (see Results section for more details), thus preserving *scDblFinder*’s top-ranking overall performance. However, it is worth noting that no single method produced the best results across all datasets, thus highlighting the need to test and benchmark methods across several datasets and suggesting that some strategies might have advantages and disadvantages depending on the situation.

## Methods

### scDblFinderPy implementation

To ensure inter-operability, *scDblFinderPy* was built directly on the *AnnData/Scanpy* ecosystem (Virshup et al., 2021; Wolf et al., 2018). It operates on AnnData objects, requiring raw count data in *adata*.*X* or an *adata*.*layers[‘counts’]* layer, and returns per-cell doublet scores and labels in *adata*.*obs*. Numerical routines are implemented with *NumPy, SciPy* and *pandas* (Bressert, 2012; McKinney et al., 2011), multiple-testing correction for marker-based feature selection was implemented using *statsmodels* (Seabold, Perktold, et al., 2010), and k-means and exact nearest-neighbour search fall back to *scikit-learn* (Pedregosa et al., 2011) on CPU. Where R’s *scDblFinder* calls *Bioconductor* packages with no direct Python equivalent, we substitute the closest widely-used Scanpy/Python counterpart: *BiocNeighbors::findKNN* is replaced by *Scanpy’s pp*.*neighbors* on CPU and *rapids-singlecell’s* neighbours function on GPU, and bluster/igraph-based Louvain clustering is replaced by the *igraph* Python bindings (`community_multilevel`), which run the same underlying algorithm that *igraph::cluster_louvain* calls in R (Blondel et al., 2008; Csardi et al., 2013). Gradient-boosted classification uses XGBoost (Chen and Guestrin, 2016) via its Python API — the same underlying library the R package already calls through its own bindings, so this is a change of interface rather than of the algorithm itself.

Fidelity to the original method was the primary goal when developing this package, and *scD-blFinderPy’s* pipeline structure closely mirrors R’s *scDblFinder* step for step. However, there were several instances where decisions needed to be made in order to navigate around the different implementations of certain libraries and modules across the two programming languages. One such decision involved which k-means algorithm to use: while the default algorithm in R is Hartigan-Wong’s, *scikit-learn* (Pedregosa et al., 2011) only implements Lloyd’s, which was therefore used. Another example would be the gene ranking used ahead of the preclustering PCA: in the R *scDblFinder* package, *fastcluster* uses *scater::runPCA(ntop=*…), which ranks genes by raw variance, as opposed to *Scanpy’s* default dispersion based gene selection. Since the resulting output from this process is used in several downstream steps and any deviations could significantly impact the resulting accuracy, we have decided to replicate this raw variance ranking in our Python package instead of relying on the default *Scanpy* implementation.

### GPU acceleration

GPU acceleration is provided as an opt-in path using the *rapids-singlecell* package (Dicks et al., 2026), which exposes the same API as *Scanpy* (Wolf et al., 2018) and is backed by *cuML/cuGraph* (Raschka et al., 2020), thus allowing the pipeline’s preprocessing calls to be dispatched to CPU or GPU through a single backend-selection function whilst avoiding code duplication.

Count normalisation, log-transformation and PCA in both the preclustering step and the main real + artificial embedding step are run via *rapids-singlecell/cuML*. The k-means preclustering step uses *KMeans* from the *cuML* package, explicitly configured with exact *k-means++* initialisation rather than the default scalable/approximate variant so that the GPU and CPU paths solve the same clustering problem. The k-nearest-neighbour graph construction for the KNN-derived doublet features uses the GPU neighbour search from *rapids-singlecell*. The CXDS coexpression scoring step’s sparse matrix products and per-cell score aggregation are accelerated directly with *CuPy* (Nishino and Loomis, 2017), and the XGBoost training itself runs with *device=‘cuda’* and *tree_method=‘hist’* fixed on both CPU and GPU so that the two devices fit structurally identical trees rather than diverging classifiers.

Steps that remain CPU-bound are those that are either inherently small-scale or not accelerator-friendly, such as Louvain community detection (Blondel et al., 2008), which always runs on the deliberately small meta-cell graph via *igraph* (Csardi et al., 2013), artificial-doublet generation, marker/feature selection, and final threshold optimisation, all of which are *pandas/SciPy*-driven (Bressert, 2012; McKinney et al., 2011) and therefore not offloaded.

GPU mode can be enabled with a single boolean argument, namely *use_gpu*, which is passed to the *compute_doublet_score()* function (the default is set to False). Note that if *rapids-singlecell* is not importable in the active environment, the pipeline emits a warning and transparently falls back to the CPU backend instead of failing.

### Package versions

We used the following package versions for the methods compared: *pyscDblFinder* 0.2.1 (downloaded at https://github.com/omicverse/py-scDblFinder); *DoubletFinder* 2.0.6; *scDblFinder* 1.24.10; *scds* 1.26.1; *scDblFinderPy* 0.1.0; *Vaeda* 0.0.30; *Scrublet* 0.2.3.

## Results

### State-of-the-art accuracy in doublet detection

In the original *scDblFinder* paper, we compared the R package with an independent benchmark developed by Xi and Li, 2021a and noted that *scDblFinder* had the highest mean area under the precision-recall (PR) curve and ranked first in the majority of the datasets (Germain et al., 2022). Here, we reproduced a similar benchmark, where we used the latest versions of all packages, included both CPU and GPU results for our Python based implementation, and also included results from an additional package, Vaeda (Schriever and Kostka, 2023). During development, another group also reimplemented *scDblFinder* in python, and we therefore also included it (*pyscDblFinder*).

The results shown in Figure 1 clearly show that both Python implementations mimic the performance of their R counterpart. This being said, no individual method achieved the highest accuracy across all datasets. *scDblFinderPy* was superior to *pyscDblFinder* on a majority (10/16) of datasets, although the difference is not statistically significant and overall very mild.

**Figure 1.**
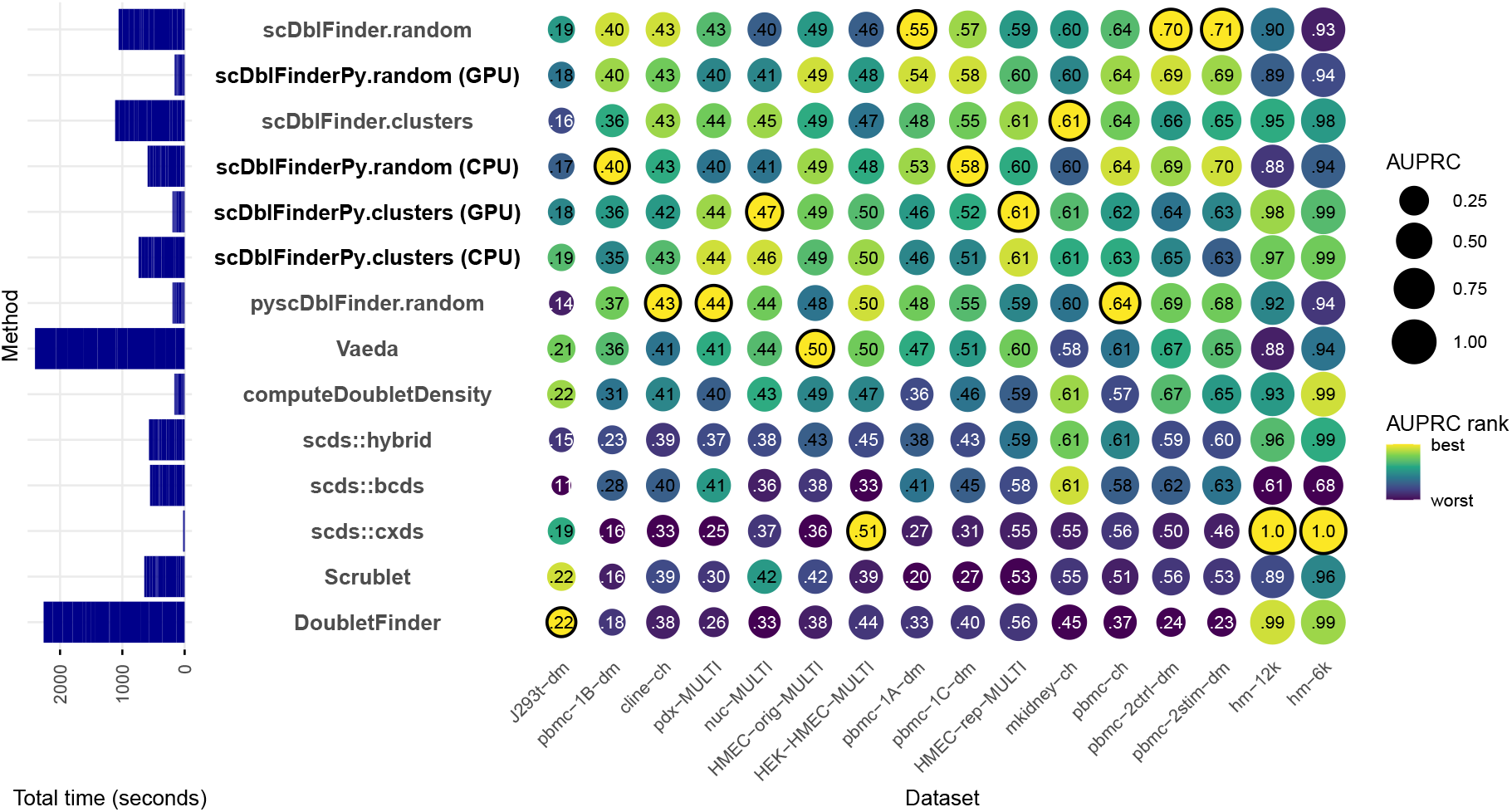
Accuracy (area under the precision and recall curve; AUPRC) of doublet identification using alternative methods across the 16 benchmark datasets from Xi and Li, 2021a. The size of the dots indicates the relative ranking for the dataset, and the numbers indicate the actual area under the (PR) curve. The AUPRC values are averaged across 3 random seeds. For each dataset, the top method is circled in black. The methods are ranked according to mean AUPRC, and the datasets are ranked by maximum AUPRC. Methods newly developed in the context of the present study are in bold.

### Computational speedup

Another benefit of the original *scDblFinder* package was its computational efficiency, and we therefore benchmarked it against our *scDblFinderPy* implementation, comparing it both in a CPU-only setting as well as in a GPU-enabled setting. On a single CPU, the Python re-implementation achieved a nearly 2-fold speedup compared to its R counterpart (Figure 1, left), which was further increased when GPU access was enabled. Notably, however, *pyscDblFinder* achieved a similar speed with CPU only.

### Mild gain from averaging across runs

With the exception of *cxds*, doublet calling methods involve the generation of random doublets, such that calling e.g. *scDblFinder* twice on the same data leads to highly-correlated results, but for some cells the doublet score can vary strongly due to the randomly-generated doublets. While generating more doublets can mitigate this problem, a more sensible approach is to run the method twice, and average the results. For many methods this is computationally more economical (because computational complexity does not increase linearly with the number of cells) and, for *scDblFinder*, more accurately reflects the amplification of random initial biases through the iterative procedure. We therefore tested to what extent accuracy can be gained by running the methods multiple times (Figure 2). While there was a clear gain for most methods, it was very mild, especially for *scDblFinder*-based methods.

**Figure 2.**
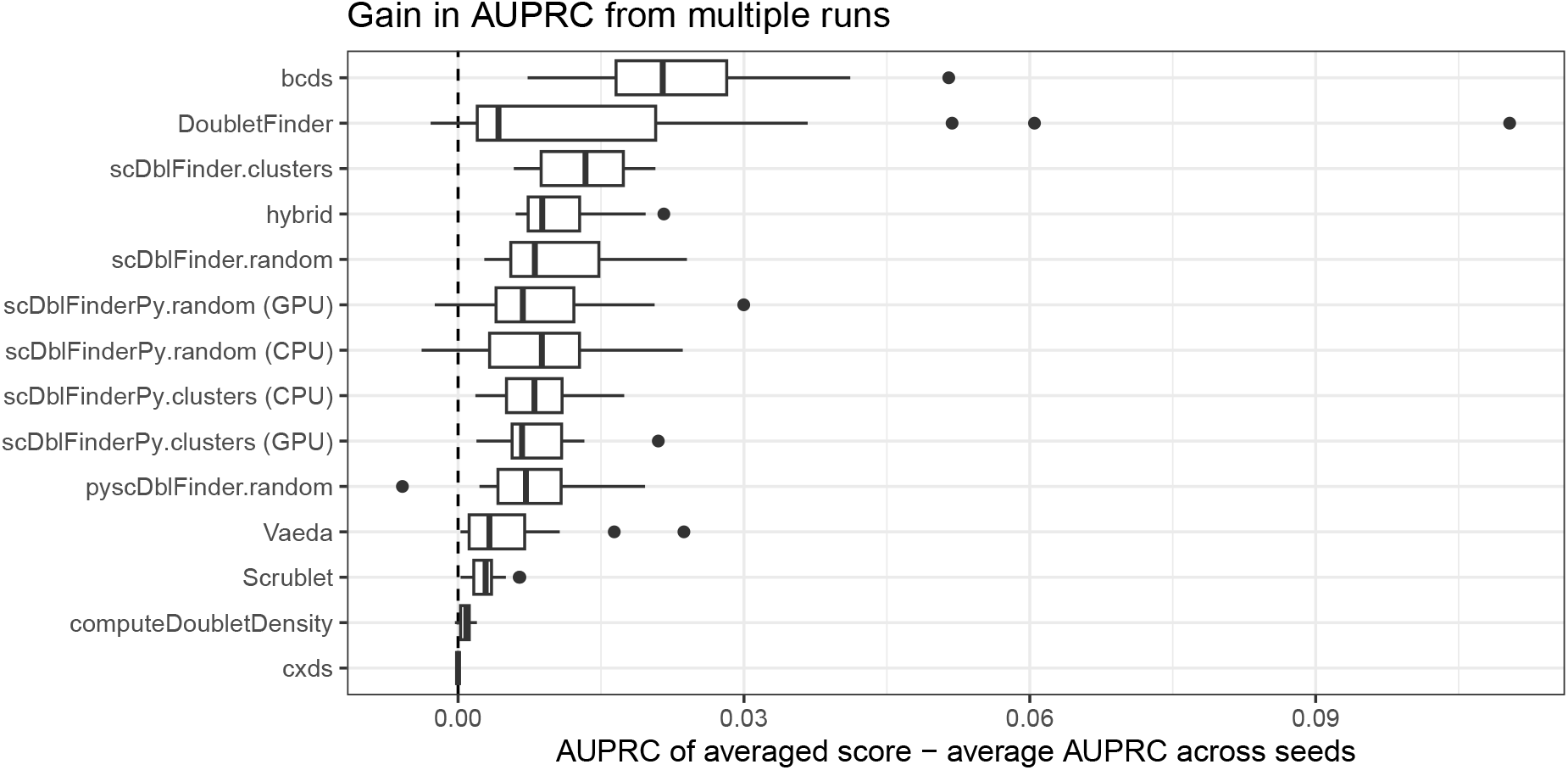
Average difference in Area Under the Precision-Recall Curve (AUPRC) computed on the cell-wise average of doublet scores across random seeds and the average of the AUPRC computed on individual random seeds. The dots represent the 16 benchmark datasets, and a positive value indicates a gain from averaging scores across runs/seeds.

## Conclusion

*scDblFinderPy* offers a Python re-implementation of the core *scDblFinder* R method to facilitate adoption to the Python community while preserving the original R package’s mixed doublet-generation strategy and providing robust and accurate results across the same benchmark datasets. This Python based approach, similar to the R method, also offers additional gains in speed and scalability through optional GPU acceleration via the *rapids-singlecell* package. As with *scDblFinder*, doublet scores remain directly interpretable as probabilities, and the trade-off thresholding procedure is preserved, thus keeping the original method’s usability intact across languages, although the results will not be (and could not even be made to be) exactly the same. During the course of the study, a similar re-implementation (*pyscDblFinder*) was done by another group, showing similar performance. In conclusion, these packages port *scDblFinder*’s state-of-the-art performance to the Python-based community.

## Funding

The authors declare that they have received no specific funding for this study.

## Conflict of interest disclosure

The authors declare that they comply with the PCI rule of having no financial conflicts of interest in relation to the content of the article.

## Data & Software availability

The source code for the *scDblFinderPy* package is available from https://github.com/ETHZ-INS/scDblFinderPy. The software is released under the GNU Public License (GPL-3). Additionally, the code to reproduce the analyses and figures is available from https://github.com/ETHZ-INS/scDblFinderPy_paper.

## Notes

### Competing Interest Statement

The authors have declared no competing interest.

